# Genetic and environmental perturbations lead to regulatory decoherence

**DOI:** 10.1101/369306

**Authors:** Amanda Lea, Meena Subramaniam, Arthur Ko, Terho Lehtimäki, Emma Raitoharju, Mika Kähönen, Ilkka Seppälä, Nina Mononen, Olli T. Raitakari, Mika Ala-Korpela, Päivi Pajukanta, Noah A. Zaitlen, Julien F. Ayroles

## Abstract

Correlation among traits is a fundamental feature of biological systems. From morphological characters, to transcriptional or metabolic networks, the correlations we routinely observe between traits reflect a shared regulation that remains poorly understood and difficult to study. To address this problem, we developed a new and flexible approach that allows us to identify factors associated with variation in correlation between individuals. Here, we use data from three large human cohorts to study the effects of genetic variation and environmental perturbation on correlations among mRNA transcripts and among NMR metabolites. We first show that environmental exposures (namely, infection and disease) lead to a systematic loss of correlation, which we define as ‘decoherence’. Using longitudinal data, we show that decoherent metabolites are better predictors of whether someone will develop metabolic syndrome than metabolites commonly used as biomarkers of this disease. Finally, we show that correlation itself is a trait under genetic control: specifically, we mapped and replicated hundreds of ‘correlation QTLs’, which often involve transcription factors or their known target genes. Together, this work furthers our understanding of how and why coordinated biological processes break down, and highlights the role of decoherence in disease emergence.

## Introduction

A major goal in evolutionary and medical genetics is to identify the coordinated regulatory processes that differ between individuals as a function of their disease state, environment, or genetic background. One powerful approach for doing so is to identify the effect of environmental or genetic perturbations on the degree to which genes are correlated at the mRNA level – a phenomenon known as ‘co-expression’. However, we still have a poor understanding of how genetic variation can modify correlations between genes, or how environmental perturbations alter essential patterns of co-regulation. From a medical perspective, identifying the factors associated with changes in co-expression between healthy and sick individuals should point toward key regulatory changes driving phenotypic differences between groups^1–5^ (and thus allow us to identify potential biomarkers of disease). From an evolutionary perspective, work on decanalization^6–8^ indicates that gene regulatory networks evolve over many generations under stabilizing selection, and new mutations or novel environments may disrupt fine-tuned connections and breakdown co-regulation. Decanalization has been advanced as one of the most compelling explanations for the recent rise of non-communicable diseases in humans^7,9^, with the hypothesis being that major shifts in diet, lifestyle, or pathogen exposure lead to dysregulation of regulatory programs that evolved under different environmental conditions. However testing this hypothesis remains challenging in practice.

Our ability to move forward in understanding how environmental and genetic perturbations affect molecular correlations is limited by the available methodology (reviewed in^5,10,11^). In particular, almost all studies of differential co-expression to date have relied on two types of approaches: (i) building co-expression networks separately within each group of interest (e.g., diseased versus healthy) and contrasting the two with a network comparison tool; or (ii) asking whether predefined sets of genes are differentially co-expressed between groups. Almost universally, these approaches are only designed for comparisons between two groups, and none can accommodate continuous predictor variables. Further, these approaches are only designed for data sets in which no confounders or covariates may bias co-expression estimates, which could lead to false conclusions if not accounted for^12^. This limited flexibility has made it difficult to identify individual-specific factors that predict co-expression in humans or other organism that are not amenable to controlled manipulations.

To address this gap, we present a novel approach for asking whether the degree of correlation between two traits is predicted by a variable of interest. Importantly, it relies on the flexible linear modeling framework and can thus accommodate covariates and continuous predictor variables. Our approach is based on the fact that the correlation between two traits within a sample is equal to the average product of the two traits across individuals, after the data are mean centered and scaled. By extension, to obtain a measure of the degree of correlation between two traits for each individual in a sample, we can simply take the vector of products between the two traits after normalization. We call this approach ‘*Correlation by Individual Level Product’* (CILP), because for each individual a product can be estimated and used to model correlation as we would any other continuous outcome variable (Figure 1).

**Figure 1.**
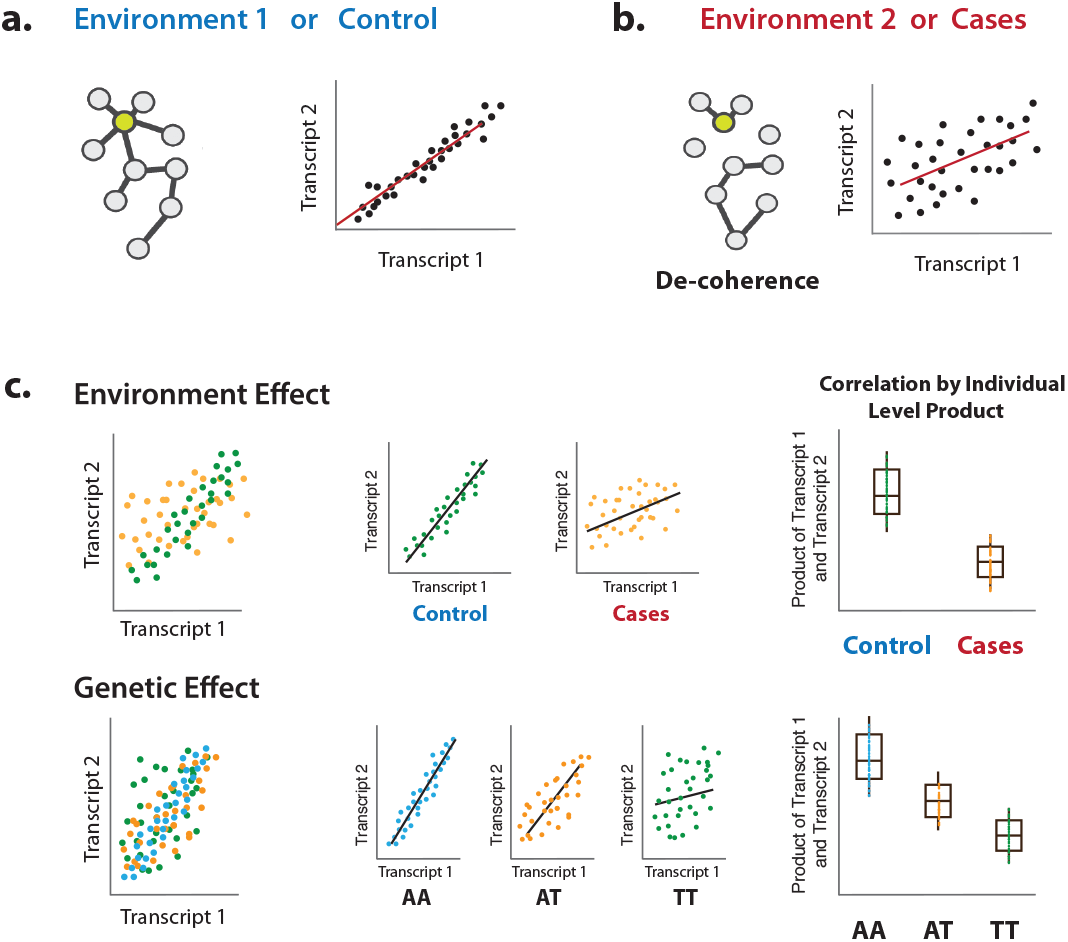
Illustration of decoherence and ‘Correlation by Individual Level Product’ (CILP). (a) Co-expression network in two different environments. Environment 1 represent normal (or control/baseline) conditions where the expression levels of gene 1 and gene 2 are highly correlated (co-regulated). (b) Environment 2 represents a stressful (or unhealthy) condition leading to lower correlation between the expression levels of gene 1 and gene 2. At the network level, this translates into a lower network degree (lower average correlation across genes or fewer connected nodes). We call this change in the correlation structure between genes ‘decoherence’. This is what we could expect if stressful conditions lower transcriptional robustness and lead to dysregulation of gene expression. (c) Difference in correlation between the expression levels of gene 1 and gene 2 in cases versus controls or between genotypes, which translates into an average difference in CILP between these groups.

Leveraging this approach, we explore the effects of environmental and genetic variation on the degree to which two molecular traits are correlated. First, we test the hypothesis that stressful physiological conditions (here, bacterial infection or metabolic syndrome) lead to a loss of correlation among molecular traits that are correlated under normal conditions. We refer to this loss of correlation in an infected or diseased state, relative to a baseline or healthy state, as ‘decoherence’. We test for decoherence using gene expression data derived from monocytes at baseline or following stimulation with lipopolysaccharide^13^, a component of bacterial cells walls and a potent stimulant of the innate immune response^13^. We also test for this loss of regulatory homeostasis using blood-derived NMR metabolite data from the Young Finns Study (YFS)^14^. Strikingly, we find strong evidence for decoherence in both data sets. Finally, we use CILP to map ‘correlation QTL’, defined as SNPs that affect the magnitude of the correlation between two mRNA transcripts. Using genotype and whole blood-derived gene expression data from the Netherlands Study of Depression and Anxiety^15^, we identity and replicate hundreds of correlation QTLs. Together, our new approach allows us to identify genetic variant and environmental factors that disrupt molecular co-regulation. Further, the flexible, robust approach we propose opens the door to future investigations of the causes and consequences of trait covariation in many contexts.

## Results

### Using Correlation by Individual Level Product (CILP) to test for sources of variance in correlation

Let *y*_1_,*y*_2_ be two outcomes measured across a population sample, with means *y̅*_1_, *y̅*_2_ and variances 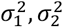, respectively. We wish to associate the correlateion between these two variables: 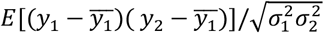 with some random variable *x*. We propose the following statistical test. First, we calculate the demeaned product of the outcomes: [(*y*_1_– *y̅*_1_ (*y*_2_– *y̅*_1_)], and then normalize by the square root of the product of variances: 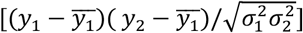. The resulting vector of values represents the estimate of the correlation within each individual in the sample, which can subsequently be modeled using approaches appropriate for continuous outcome variables. In practice, we use linear regression or linear mixed effects models to test for associations between a given set of products and *x* (controlling for covariates).

### Simulations reveal power to detect sources of variance in correlation across many scenarios

To confirm that our approach does not result in biased p-value distributions, and to understand the power of CILP across a range of effect sizes and sample sizes, we first performed extensive simulations. We focused our simulations on the identification of ‘correlation QTL’, defined as SNPs that affect the magnitude of the correlation between two mRNA transcripts (however, we note that the results are generalizable other types of predictor variables). In each case, we simulated 10,000 pairs of genes, such that samples originating from different groups (i.e., different genotypic classes) exhibited different levels of correlation for each gene pair. We did so using the multivariate normal distribution to simulate pairs of continuous distributions (but see Supplementary Figure 1 for results where count data were simulated from a negative binomial distribution). Following simulation, we CILP and linear models to detect differences in correlation as a function of group membership.

With an effect size of 0.3 and n=1000, we detected 98.19% of simulated true positive correlation QTLs at a threshold of p=0.05, and 56.57% of true positives after correcting for multiple testing (using Bonferroni correction; Supplementary Figure 1). Under the null, where the effect size was set to zero, we detect 4.54% of correlation QTLs at a threshold of p=0.05 and after no correlation QTL after correcting for multiple testing (Supplementary Figure 1). To ensure that changes in the mean and variance of the traits of interest do not increase our false positive rate, we assessed our power to detect correlation QTLs when: (i) the focal SNP also affected the mean expression levels of one or both genes (i.e., it was an eQTL) and (ii) the focal SNP also affected the variance in expression levels for one or both genes (i.e., it was a varQTL). Neither the presence of eQTLs nor varQTLs increased the proportion of false positive correlation QTL detected. As expected, the presence of eQTLs nor varQTLs increased the proportion of false positive correlation QTL detected. As expected, the presence of strong varQTLs did decrease power to detect correlation QTLs (Table 1). We also found that including the expression levels of both genes as covariates in our models did not affect our power to detect correlation QTLs (Supplementary Figure 1).

**Table 1.** Power to detect simulated correlation QTL when the focal genetic variant also affects the mean or variance of one or both of the correlated genes (n=1000 for all simulations).

Using Correlation by Individual Level Product (CILP) to test for sources of variance in correlation
| Simulation parameters |  |  |  |  |  |  |  |  |
| --- | --- | --- | --- | --- | --- | --- | --- | --- |
| Correlation QTL effect<sup>1</sup> | x | x | x | x | x | x | x | x |
| eQTL effect<sup>2</sup> (gene 1) |  | x | x |  |  | x |  |  |
| eQTL effect<sup>2</sup> (gene 2) |  |  | x |  |  |  |  | x |
| vQTL effect<sup>3</sup> (gene 1) |  |  |  | x | x | x | x | x |
| vQTL effect<sup>3</sup> (gene 2) |  |  |  |  | x |  |  |  |
| Results |  |  |  |  |  |  |  |  |
| Power (proportion of true positives with p<0.05) | 0.9999 | 1 | 0.9998 | 0.3777 | 0.0795 | 0.3858 | 0.2901 | 0.2932 |
| False positive rate (proportion of true negatives with p<0.05) | 0.0527 | 0.0488 | 0.0495 | 0.0472 | 0.0492 | 0.0507 | 0.0501 | 0.0501 |

### Immune challenges results in decreased co-regulation of gene expression

Next, we used gene expression data collected from human monocytes, to ask whether patterns of co-expression differed between cells assayed at baseline versus following treatment with LPS for 2 or 24 hours (n=214 samples were assayed at baseline, as well as 2 and 24 hours post LPS stimulation). We focused our analyses on 1460 genes that were differentially expressed at the 2-hour time point (FDR<5%), and tested for differences in correlation between unexposed and LPS-exposed cells across 1063611 possible transcript pairs (equivalent to 1460 chose 2). We found 958 gene pairs with a significant change in correlation between the two conditions (uninfected/baseline versus 2 hours of LPS stimulation), 461 of which replicated at this threshold with similar effect sizes at the 24-hour time point (note that 52 genes pairs replicated at a Bonferroni-corrected p<0.05; correlation between effect sizes at the two time points: R^2^=0.12, p<10^-16^; binomial test for concordance of effect size direction: p<10^-16^; Figure 2). We observed no relationship between the magnitude of the difference in mean expression levels between conditions and the magnitude of the difference in correlation between conditions, suggesting our results are not driven by statistical artifacts associated with large mean changes in gene expression following LPS stimulation (Supplementary Figure 2).

**Figure 2.**
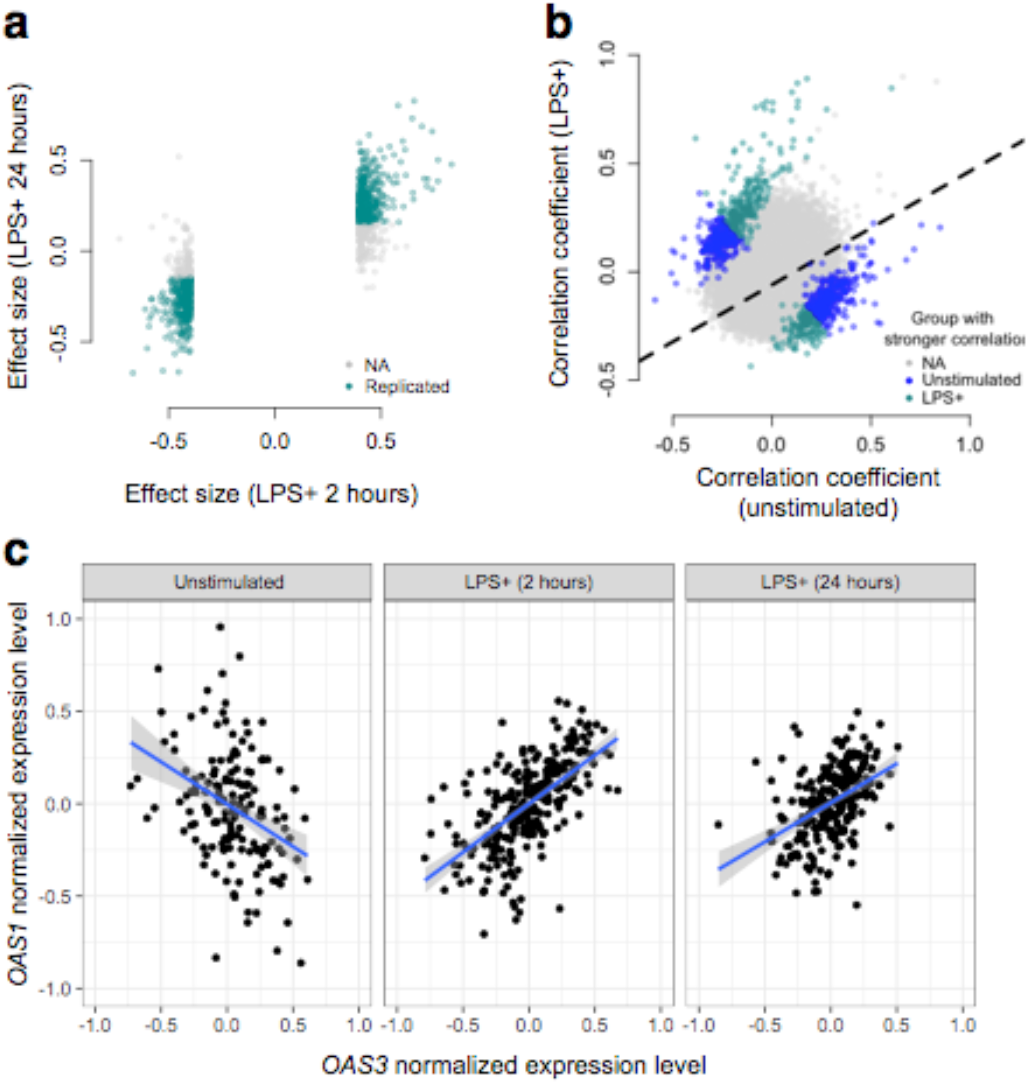
Simulated bacterial infection (i.e., treatment with LPS) leads to decoherence in primary monocyte gene expression. **(a)** Correlation changes have similar effect sizes in the 2-hour LPS stimulation and the 24-hour LPS stimulation. **(b)** Gene expression values for OAS1 and OAS3 at the baseline, LPS 2-hour, and LPS 24-hour conditions show higher correlation after LPS stimulation. **(c)** Expression values for PPBP and GNG11 indicate lower correlation after LPS stimulation. **(d)** Density plots for pairwise correlations at baseline, 2 hours after LPS stimulation, and 24 hours LPS stimulation show that there is a shift towards less correlation upon stimulation.

In total, we identified and replicated gene pairs that are more highly correlated in cells assayed 2 hours post LPS stimulation versus at baseline, as well as gene pairs that lose correlation during bacterial infection. For example, *OAS1* and *OAS3* – two key genes in the type I interferon pathway^16^ – are more strongly positively correlated in the LPS condition relative to baseline (at both 2 and 24hrs post stimulation, p=1.19×10^-7^ and p=1.20×10^-23^, respectively; Figure 2). Overall, however, we found greater support for the opposite pattern of correlation change: 61% (at the 2 hour time point) and 73% (at the 24 hour time point) of significant transcript pairs were more strongly correlated across individuals in uninfected versus LPS stimulated cells. Importantly, this represents a significant bias toward a loss of correlation (i.e. decoherence) following two hours of bacterial infection (p=1.05×10^-7^, log_2_ odds=0.503, Fisher’s exact test), with an even stronger bias toward dysregulation after 24 hours of immune challenge (p<10^-16^, log_2_ odds=0.941, Fisher’s exact test).

### Metabolic syndrome disrupts metabolite co-regulation

#### Identification of disease-associated variation in metabolite correlation

We next applied our correlation test to whole blood-derived NMR metabolite data collected from a population-based, longitudinal study of young Finnish individuals (the cardiovascular risk in young Finns study, abbreviated ‘YFS’)^14,17^. Our dataset included 159 metabolite measures, and, after filtering (see Methods), we retained 11491 metabolite pairs for analysis. These metabolites were measured across three time points (2001: n=1564, 2007: n=1498, 2011: n=1501); however, our analyses did not focus on changes in correlation across time, but rather, changes in correlation between individuals that were healthy versus those that met the criteria for metabolic syndrome^18^.

To test the hypothesis that disease disrupts homeostasis and perturbs molecular co-regulation, we asked whether specific pairs of metabolites were correlated in healthy individuals, but no longer correlated in those with metabolic syndrome. Across 11491 unique metabolite pairs, the estimate of the correlation coefficient in healthy people and in those with metabolic syndrome tended to be similar (R^2^=0.62, p<10^-16^; linear model controlling for age; Figure 3). However, for a subset of metabolite pairs, we found strong effects of health status on the magnitude of the correlation: 1528 (74.4%) metabolite pairs were more correlated in healthy individuals relative to those with metabolic syndrome, and 619 (25.6%) metabolite pairs showed the opposite pattern (FDR<5%; linear mixed model controlling for age, sex, year, and individual identity). This represents a 2.20x enrichment of metabolite pairs that appear to become dysregulated (i.e., that lose correlation) following the onset of disease relative to chance expectations (p<10^-16^, Fisher’s exact test). Importantly, this set of 1528 metabolite pairs that become dysregulated includes metabolites that are both more highly expressed (n=93 metabolites, FDR<5%) and more lowly expressed (n=32) in individuals with metabolic syndrome, as well as metabolites that do not significantly differ between sample groups (n=4).

**Figure 3.**
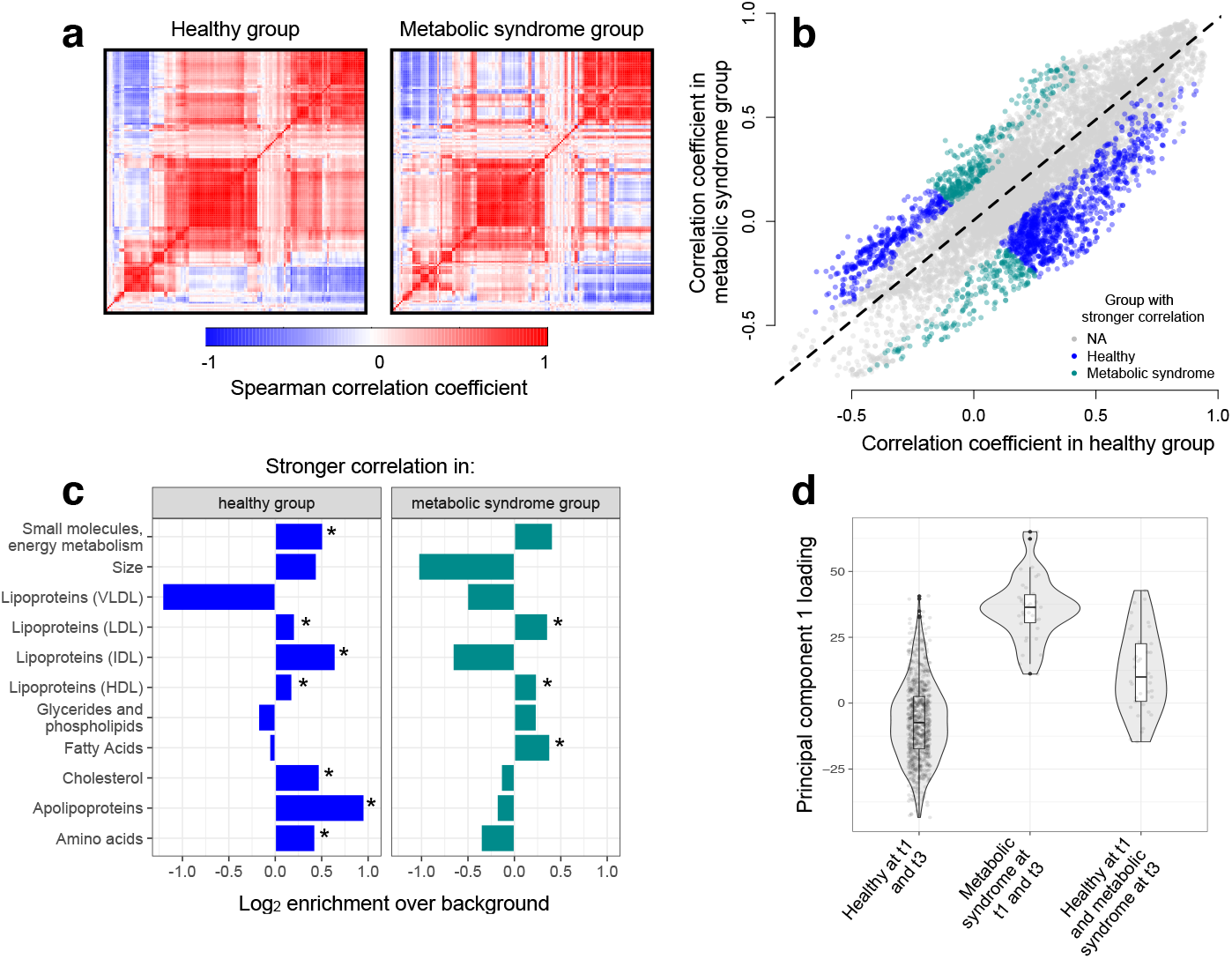
Metabolic syndrome leads to decoherence among particular metabolite pairs. (**a**) Correlation matrices showing the magnitude of the Spearman correlation coefficient, for healthy individuals and those with metabolic syndrome. (**b**) Comparison of the Spearman correlation coefficient estimated in healthy individuals (x-axis) versus those with metabolic syndrome (y-axis). Overall, the magnitude of the correlation is similar between groups (R^2^=0.62, p<10^-16^); however, for a subset of metabolite pairs, we detect stronger correlations in the healthy (blue dots) or metabolic syndrome class (green dots). (**c**) Categorical enrichment of metabolites that exhibit stronger pairwise correlations in the healthy or metabolic syndrome class (x-axis: log_2_ odds ratio from a Fisher’s exact test; y-axis: 11 functional classes tested (annotations taken from^19^). Asterisks indicate significant enrichment. (**d**) A composite measure of metabolites that are strongly decoherent at the first time point (i.e., that show decreases in correlation in individuals with metabolic syndrome) can predict an individual’s future health status. Y-axis: Principal component 1 of 34 metabolites, measured at the first time point, with the strongest evidence for dysregulation. X-axis: values are stratified by whether an individual was healthy at the first (t0) and the last (t3) time point, had metabolic syndrome at t0 and t3, or developed metabolic syndrome between t0 and t3 (linear model, p=3.03×10^-10^).

#### Metabolic syndrome affects co-regulation of particular metabolite classes

We found overall support for the hypothesis that metabolic syndrome disrupts the correlation structure that exists in healthy individuals. To identify the specific processes targeted by metabolic syndrome, we assigned each metabolite in our dataset to one of 11 functional classes (as in^19^; Supplementary Table 2), and asked whether each class was enriched for metabolites that exhibited decreases in correlation strength. Here, we found the strongest enrichment for apolipoproteins (hypergeometric test, odds ratio=1.94, p=2.34×10^-13^), measures of total cholesterol (odds ratio=1.39, p=1.58×10^-15^), and small molecules involved in energy metabolism (odds ratio=1.43, p=2.75×10^-13^). Among metabolite pairs that showed the opposite pattern (i.e., were more strongly correlated in individuals with metabolic syndrome), we found an enrichment of fatty acids (odds ratio=1.30, p=7.10×10^-6^), as well as HDL (odds ratio=1.18, p=1.53×10^-5^) and LDL (odds ratio=1.28, p=3.58×10^-6^) lipoproteins (Figure 3). Together, these results suggest that metabolite co-regulation is strongly perturbed by disease, and further, that particular classes of metabolites are more sensitive and prone to dysregulation than others.

Finally, because our dataset included metabolite data collected from the same individuals across multiple time points, we asked whether metabolite pairs that became dysregulated (i.e., lost correlation) following the onset of disease could be used to predict which individuals would develop metabolic syndrome at a later time point. To do so, we performed PCA on the 34 metabolites that displayed the strongest evidence for dysregulation at the first time point, and used the first principal component to predict an individual’s health status at the last time point (see Methods). Strikingly, we found that this measure was predictive of whether an individual would develop metabolic syndrome (R^2^=0.056, p=3.03×10^-10^, AIC=-448.40; Figure 3), more so than triglyceride levels at the first time point (a classic biomarker of metabolic syndrome; R^2^=0.046, p=6.32×10^-9^, AIC-432.61) or than an index created from the 34 metabolites with the strongest mean differences between healthy and metabolic syndrome individuals at the first time point (R^2^=0.037, p=1.42×10^-6^, AIC=-422.08).

### Genetic variation impacts co-expression of metabolism-related genes

#### Detection and replication of hundreds of correlation QTL

Next, we used our correlation test to identify genetic variants that control the degree of correlation between a pair of gene transcripts (a pattern we refer to as ‘correlation QTL’). Genotypic effects on co-expression could arise through several possible mechanisms. For example, a SNP that disrupts a transcription factor (TF) binding site in a promoter would lead to low levels of co-expression between the TF and the target gene, but only for individuals carrying the disrupting variant. In these individuals, increased expression of the TF would fail to increase expression of the target gene, leading to an association between SNP genotype and variation in co-expression. This is one mechanistic scenario that has been repeatedly proposed to generate variation in co-expression^5,20^, with some empirical support^19^. However, the relationship between genetic variation and co-expression is almost entirely unexplored, suggesting that alterative mechanisms may exist that have yet to be uncovered.

To map SNPs that affect the magnitude or direction of pairwise gene expression correlations, we applied our correlation test to genotype and whole blood-derived gene expression data from the Netherlands Study of Depression and Anxiety (abbreviated ‘NESDA’ ^15^). We used a filtered set of 93,197 SNP genotypes and 33,302 gene expression measurements collected for n=2,477 individuals in our discovery dataset and n=1,337 individuals in our replication dataset (see Methods). Because our dataset was not well-powered to test all possible pairwise combinations of gene expression measurements against all SNPs, and because we were interested in understanding co-expression patterns among genes important to metabolic diseases, we focused on the 475 probes most strongly associated with BMI. In total, we identified 484 associations between a SNP and variation in co-expression in our NESDA discovery dataset (at a 10% FDR threshold, corresponding to p<4.6×10^-9^; linear model controlling for sex, age, smoking behavior, major depressive disorder, red blood cell counts, year of sample collection, study phase, and the first 5 principal components from a PCA on the filtered genotype call set). These 484 associations involved 247 unique probe sets and 51 genotyped SNP, each of which was involved in 1-424 (mean ± s.d. = 3.91 ± 26.9) and 1-173 (mean ± s.d. = 9.43 ± 34.1) correlation QTL, respectively (Figure 4; Supplementary Table 3).

**Figure 4.**
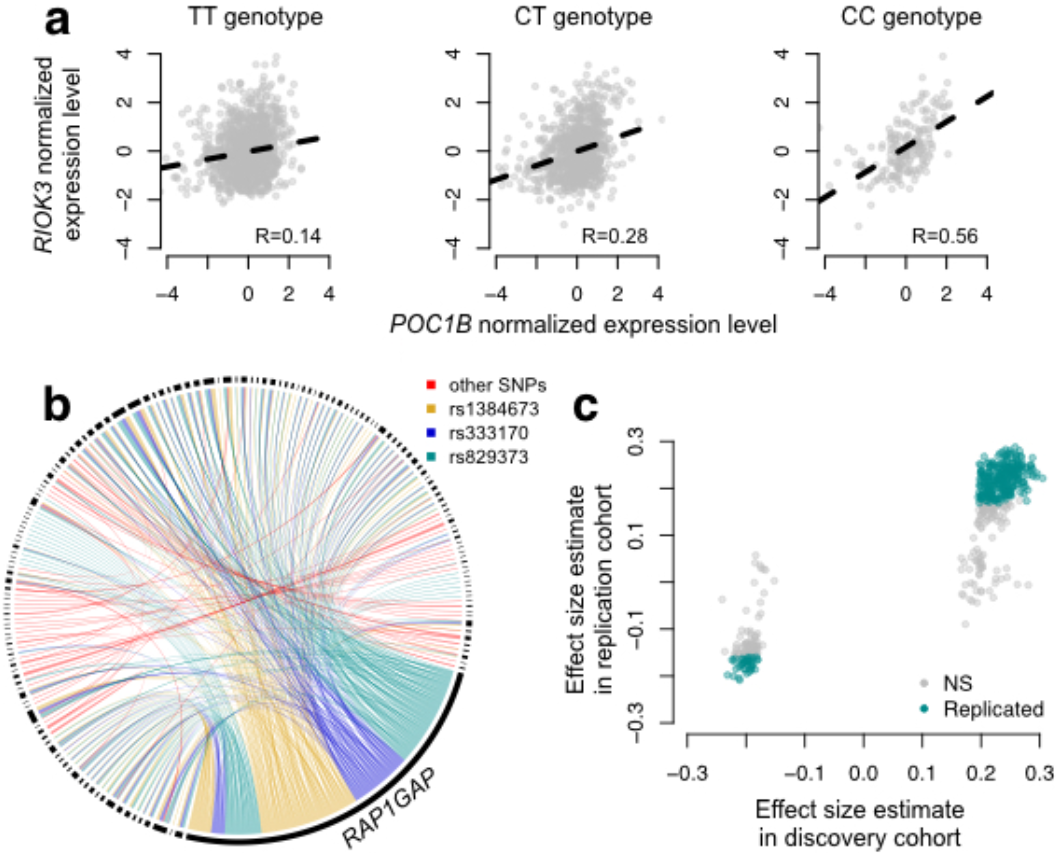
CILP approach reveals hundreds of correlation QTLs. (**a**) Example of a correlation QTL, where the SNP rs10953329 controls the magnitude of the correlation between the mRNA expression levels of *POC1B* and *RIOK3.* (**b**) Gene pairs involved in significant (FDR<10%) correlation QTL. Each black segment represents a gene, and each line connecting two segments represents a significant correlation QTL. Lines are colored by the identity of the SNP controlling the magnitude of the correlation between the gene pair. (**c**) Many correlation QTL identified in the NESDA discovery set (n=2477) replicate in a second set of NESDA participants (n=1337). Plot shows effect sizes for each correlation QTL, estimated in the discovery or replication cohort (effect sizes are derived from in *matrixEQTL*^50^). Points are colored to indicate whether a given correlation QTL passed Bonferroni correction in the replication dataset.

To confirm our results, and overcome any potential biases raised by sub-selection of the data, we replicated our correlation QTL in a separate set of NESDA participants. Here, we found that 304/484 correlation QTL replicated (at a Bonferroni corrected p=1.03×10^-4^, n=1337; 428/484 correlation QTL replicated at a 10% FDR; Figure 4). Further, in whole-blood derived gene expression data from 1414 YFS participants, we found that 47/74 testable correlation QTL replicated at a 10% FDR (0 replicated at a Bonferroni corrected p=6.76×10^-4^, but the direction of the effects consistently agreed across datasets (binomial test, p<10^-16^); see Methods).

#### Correlation QTLs reveal transcription factor biology

For the list of 484 correlation QTLs we identified, we performed several follow-up analyses to gain biological and mechanistic insight. First, we asked whether the set of genes involved in significant correlation QTLs were enriched for (i) particular biological processes and pathways, (ii) known TFs, or (iii) known targets of TFs compared to the background set of all genes tested for correlation QTLs (using publicly available databases of gene ontologies and transcription factor-gene target associations ^21,22^). We strongly expected genes involved in correlation QTLs to be enriched for TFs or known targets of TFs, given that the mechanisms that have been proposed to generate genetic effects on variation in correlation almost universally involve genotype-dependent TF activity or disruption of TF binding sites^5,20^. In support of these ideas, genes involved in significant correlation QTLs were 1.90x more likely to be TFs (hypergeometric test, p=1.7×10^-3^) and 1.52x more likely to be known targets of transcription factors (p=6.7×10^-4^) relative to the background set of all genes tested. Additionally, genes involved in significant correlation QTLs were enriched for biological processes known to be involved in metabolism and metabolic disease^23,24^ such as cellular response to oxidative stress (hypergeometric test, odds ratio=2.21, p<10^-16^), intracellular signal transduction (log_2_ odds ratio=1.44, p<10^-16^), and mitophagy (the selective degradation of mitochondria; log_2_ odds ratio=0.82, p=0.018; Supplementary Table 4).

#### Genetic variation affects the co-regulation of RAP1 targets

Strikingly, many of the correlation QTL we uncovered involved three SNPs (rs1384673, rs333170, and rs829373), which had strong effects on the degree of co-expression between one gene, *RAP1GAP*, and an additional 105 genes. These SNPs were also strong *cis* eQTL for *RAP1GAP* (Supplementary Figure 4). The function of *RAP1GAP* is to convert the transcription factor *RAP1* – a master regulator of T and B-cell activation, cell adhesion, and neuronal differentiation – from its active GTP-bound form to its inactive GDP-bound form. Intriguingly, recent work has implicated this *RAP1GAP-*facilitated ‘switch’ (between the active and inactive forms of *RAP1*) as important in metabolic disease: mice fed a high fat diet exhibit increases in the proportion of neuronal *RAP1* that is active, and this increase is sufficient to produce multisystem effects including insulin resistance, inflammation, and obesity^25^.

Given the strong links between *RAP1* and metabolic disease, as well as the known functional relationship between *RAP1* and *RAP1GAP,* we hypothesized that BMI-associated genes involved in correlation QTL with *RAP1GAP* were directly regulated by *RAP1*. In support of this hypothesis, genes involved in correlation QTL with *RAP1GAP* tended to be closer to *RAP1* binding sites (two-sided Wilcoxon-signed rank test, p=0.04), and were more likely to be directly bound by *RAP1* (Fisher’s exact test, odds ratio=2.14, p=0.03; ChIP-seq data are from mouse embryonic fibroblasts26). Together, these data point toward the mechanistic scenario depicted in SupplementaryFigure 5: *RAP1* exists in a constitutively active or inactive form depending on the genetic variation each individual harbors near *RAP1GAP,* and intra-genotypic variation in *RAP1* activity levels is only meaningful for the low activity genotype. In contrast, for individuals with one of the high activity genotypes, the targets of *RAP1* are constitutively repressed in all individuals (Supplementary Figure 5), and increasing or decreasing *RAP1* activity levels has little effect. Together, these results help to identify an under appreciated mechanism that can generate genetic effects on gene co-expression: cases where the targets of a given TF are always ‘on’ (or ‘off’) in one genotypic class, but dynamically regulated in another.

## Discussion

Patterns of transcriptional correlation are widely considered to arise from co-regulation between genes. The analysis of co-expression has become an essential tool for the functional interpretation of transcriptional variation^5,20^, with increasing relevance for medical diagnosis^10,27,28.^ However, we still have a primitive understanding of the factors that shape correlations between genes. Specifically, how do environmental perturbations alter essential patterns of co-regulation? And, to what degree are some genotypes better than others at buffering these disruptive effects? To address these fundamental questions, we developed a novel, generalizable approach to test whether any predictor variable (e.g., environment, genotype, or another variable of interest) affects the degree of correlation between a pair of measured variables. This simple approach, which relies on calculating the product between two variables after normalization, allows us to produce an individual level estimate, rather than a summary statistic for a population sample (Figure 1). Consequently, we are able to study correlation as a *bona fide* trait, and leverage the flexibility of statistical linear modeling approaches to identify factors associated with variation in correlation between individuals.

We begin by investigating the influence of two environmental perturbations on molecular co-regulation. First, we use data from a controlled experiment, where human monocytes were challenged *ex vivo* with bacterial infection^13^. Second, we draw on a dataset of 159 blood NMR metabolites collected from a population-based, longitudinal study of Finnish individuals (the YFS study^14^), and contrast correlation patterns between healthy individuals versus those suffering from metabolic syndrome. In both instances, we find that stressful environmental exposures (infection and disease) lead to decoherence, manifested as a widespread decrease in the magnitude of pairwise correlation coefficients between mRNA transcripts or metabolites (Figure 3).

Although we expect the relationship between mRNA transcripts and metabolites to change following an environmental perturbation (e.g., both LPS stimulation and metabolic syndrome clearly affect mean gene expression or metabolite levels^13,19^), the consistent direction of effects we observe on the correlation is striking. Specifically, of all metabolite and gene transcript pairs that are significantly differentially correlated, there is a strong directional bias toward a decrease in correlation following the environmental perturbation: 61-74% of significant pairs follow this pattern. In other words, under stress, some genes and metabolites that are typically co-regulated no longer are. Similar biased loses in correlation have also been observed in miRNA pairs measured in the plasma of patients suffering from cognitive impairment versus healthy controls^29^, as well as in gene expression data collected from aging versus young mice^30^ and in a wide range of cancers^31^. However, as this pattern has generally not been tested for explicitly, the degree to which it commonly characterizes disease, aging, or environmental perturbations remains to be seen.

The loss of transcriptional robustness we observe is an intuitive extension of decanalization models^7^. Decanalization models posit that, through many generations of stabilizing selection, biological systems evolve to maintain homeostasis under a certain range of environmental (or genetic) perturbations, and changing the environment beyond this range will result in homeostatic breakdown and disease^6,7^. Here, we are potentially seeing this process play out at the molecular level, where formally co-regulated processes become decoherent following a dramatic shift in the environment. Importantly, the longitudinal nature of the YFS dataset allowed us to track the health status of the same individuals over time and test the hypothesis that decoherence at the molecular level leads to disease. In support of decanalization models, we found that variation in metabolite pairs that showed the strongest evidence for decoherence at the first-time point could be used to predict which individuals would develop metabolic syndrome at a later time point. This set of metabolites are particularly interesting from a clinical perspective, as they appear to be especially sensitive to the homeostatic breakdown associated with metabolic syndrome, and could thus potentially serve as biomarkers.

Lastly, we investigated the role of genetic variation in driving variation in gene co-expression. To date, work on differential co-expression has largely focused on contrasts between cases and controls^5,10,32^, between tissues^33,34^, or between individuals inhabiting different environments^5,35^, while the genetic basis of differential co-expression has received much less attention. Our ability to map correlation QTL directly using CILP fills this gap, and reveals that the degree of correlation between transcripts is under genetic control. Our analyses build on three recent studies that came to similar conclusions using more indirect approaches and/or much smaller sample sizes. First, Nath et al. identified highly correlated groups of mRNA transcripts and metabolites (termed ‘modules’), and performed genome-wide scans to associate specific SNPs with variation in each module^19^. In a related analysis, Lukowski et al. demonstrated that mRNA transcripts with evidence for genetic correlations were more likely to be regulated by the same expression quantitative trait loci (eQTL)^36^. Finally, van der Wijst recently leveraged single-cell mRNA-seq data from 45 individuals to build personalized co-expression networks, and used these data to identify genetic variants that predicted inter-individual variation in correlation structure^37^. Together, these studies point toward pervasive genetic control of gene co-expression, and CILP provides the tools to probe this form of regulation even further. In particular, mapping correlation QTL across different environments, and compiling a more mechanistic picture of how genetic variation affects correlation dynamics (e.g., using ChIP-seq or ATAC-seq data integrated with correlation QTL scans) are major priorities for future work.

We performed extensive simulations to show that the framework we propose can detect sources of variance in correlation across a wide range of scenarios. Critically, neither the presence of mean effects (e.g. eQTL) or variance heterogeneity (e.g. varQTL) increases the false positive rate (Table 1). With respect to statistical power, an additional benefit to this approach is that the power to detect an association between variation in correlation and any predictor variable will increase directly with sample size. This would not be the case with other currently available methods, which largely focus on comparing co-expression networks constructed in one sample group versus another^5,10^; in these cases, increasing sample size would increase the precision of the estimated correlation, but not directly the power.

While we have focused here on mapping correlation QTL and environmental effects on correlation between molecular traits, our approach could be paired with many additional statistical tools and data types. For example, given a sufficiently large dataset, this approach could be used to study how genotype x environment interactions affect correlations between traits, or to investigate genetic variation in co-morbidity for a ranges of diseases. Moving beyond genetics, this approach could be used to identify the drivers of community level correlations in ecological datasets, tradeoffs (that manifest as negative correlations) between different fitness components or life history traits, and much more. In essence, questions of how and why correlations between molecular or organism-level phenotypes vary is at the heart of many fields, and we anticipate that our approach will thus be widely applicable.

## Materials and Methods

### Simulations

To empirically evaluate the statistical tests proposed above we performed a series of experiments on simulated data sets. We first simulated genotypes for a single locus with a major and minor allele, and with a minor allele frequency of 0.5. We then simulated gene expression data (across n=1000 individuals) for 10000 sets of gene pairs with correlated gene expression levels according to the following model:

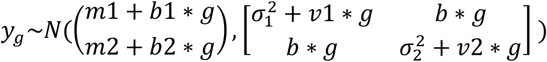

For each genotype *g* (which can take the values 0 (homozygous major), 1 (heterozygous), or 2 (homozygous minor)), we simulated gene expression values for the two focal genes with covariance *b*^*^*g*. True positives were simulated with *b* ranging from 0.05 to 0.5, and null correlation QTLs were simulated with *b* =0. The two focal genes had mean expression levels of *m1* and *m2*, respectively. *b1* and *b2* model the expression changes with respect to genotype (i.e., the effect size of the eQTL), and *v1* and *v2* model changes in variance with respect to genotype (i.e., the effect size of the varQTL).

### The cardiovascular risk in young Finns study (YFS)

#### Study subjects

We used phenotype, genotype, gene expression, and NMR metabolite data derived from a previously described study of unrelated Finnish young adults. The YFS is a longitudinal prospective cohort study initiated in 1980, with follow-ups carried out every 3 years. The study was designed to monitor cardiovascular disease risk factors in children and adolescents recruited from five regions in Finland: Helsinki, Kuopio, Oulu, Tampere, and Turku^14,17^. Our analyses focus on individuals for whom NMR metabolite data were generated from whole blood samples collected at the 2001 (n=2248), 2007 (n=2161), and 2011 (n=2040) follow-ups. Additionally, we replicate our correlation QTL findings from NESDA using paired genotype and whole blood-derived gene expression data available for a subset of individuals sampled at the 2011 follow-up (n=1414).

#### Metabolite data and metabolic syndrome classification

Metabolite concentrations were quantified from whole blood-derived serum samples collected in 2001 (n=2248), 2007 (n=2161), and 2011 (n=2040) using the procedures described in^38^. We focused on a set of 159 metabolite measures (following^19^), of which 148 were directly measured and 11 were derived (Supplementary Table 1). The 148 measures included molecules from 14 lipoprotein subclasses, two apolipoproteins, eight fatty acids, eight glycerides and phospholipids, nine cholesterols, nine amino acids, and ten small molecules (involved in glycolysis, as well as the citric acid and urea cycle). Annotations for each metabolite measurement were derived from^19^ (Supplementary Table 1). Measurements that failed quality control filters^38^ were treated as missing, and measurements of zero were set to the minimum detectable value for the particular metabolite^19^.

We removed individuals from our analyses that had type I or II diabetes, or who were reported to be on statin medications at the 2011 time point (these data were unavailable for the other two time points). For the remaining dataset, we classified individuals as healthy or having a metabolic syndrome-like phenotype at a given time point using a random forests approach implemented in the R package ‘party’^39,40^. We did so because only a subset of the five data types required to diagnose metabolic syndrome^18^ were available for all samples and time points. Specifically, fasting serum triglyceride levels, high-density lipoprotein (HDL) cholesterol levels, and blood glucose levels were available for all samples, but waist circumference and blood pressure were only available for a subset of 2011 samples (n=1414; for details on how these traits were measured, see^14,17^). We therefore trained a random forests classifier on the subset of the 2011 dataset that could be identified as having metabolic syndrome or not based on standard criteria^18^ and used the trained model to predict health status for the remaining samples. We used the follow features to generate the random forests: triglyceride levels, HDL cholesterol levels, blood glucose levels, BMI, and sex. We removed individuals from the dataset that were not confidently assigned to either class (i.e., individuals for whom the probability of assignment to either class did not exceed 2/3) leaving us with the following sample sizes: n=1564 in 2001, 1498 in 2007, and 1501 in 2011. Using data from the 2011 time point (where we have all information necessary to diagnose metabolic syndrome following the American Heart Association’s criteria^18^), we estimate that our healthy class includes 95% true positives and our metabolic syndrome class includes 84% true positives (Supplementary Table 5).

#### Genotype and gene expression data

While our analyses of YFS participants focused primarily on metabolite data, we used paired genotype and gene expression data collected from 1414 participants in 2011 to replicate correlation QTL discovered in NESDA. Whole blood was collected from each individual in PAXgene Blood RNA tubes and was used to perform genome-wide genotyping on a custom Illumina Human 670k BeadChip array and genome-wide mRNA quantification on the Illumina HumanHT-12 v4 Expression BeadChip (information on microarray experimental and quality control procedures are provided in detail elsewhere^41^).

Genotype data were filtered prior to analysis, and SNPs that met at least one of the following criteria were excluded from downstream analyses: (i) evidence that the SNP was not in Hardy-Weinberg equilibrium (p<10^-6^), (ii) data missing for >5% of all individuals, (iii) minor allele frequency <5%. Gene expression data were log2 transformed prior to analysis and filtered to remove probe sets that overlapped known SNPs, as well as those that measured lowly expressed transcripts. Additional details on genotype and gene expression quality control and filtering procedures are described elsewhere^42^.

### Testing for effects of metabolic syndrome on variation in correlation

#### Implementing the correlation test

To understand how differences in health status (‘healthy’ or exhibiting a metabolic-syndrome like phenotype) impact correlation structure, we applied our correlation test to a set of 159 NMR metabolite measures collected across three time points in the YFS. To do so, we first computed the Spearman rank correlation within each year for all pairs of metabolites (n=12561, equivalent to 159 chose 2), and excluded pairs from our analysis that were very highly correlated in any year (rho>0.9). For the remaining 11491 metabolite pairs, we computed the product after z-score transforming each metabolite measure, within year and within each set of healthy and metabolic syndrome-like individuals separately (using the ‘scale’ function in R). This normalization step is essential to ensure that mean or variance effects of year or health status on multiple metabolite measurements do not masquerade as variation in correlation.

Using the set of products computed for all filtered pairs of metabolites, we constructed linear mixed effects models in the R package ‘nlme’ ^43^. Specifically, we tested the degree to which each set of products was predicted by age and health status (healthy/metabolic syndrome-like), controlling for follow-up year (as a factor), sex, age, and individual identity as a random effect. We extracted the p-values associated with the health status effect, and corrected for multiple hypothesis testing. Importantly, we also conducted parallel analyses in which we permuted health status (metabolic syndrome/healthy) prior to (i) data normalization, correlation computation, and linear mixed model analyses or (ii) prior to linear mixed model analyses only; in both cases, our permutation results suggest that the null distribution approximates the expected uniform distribution (Supplementary Figure 5).

#### Enrichment of differentially correlated metabolite pairs

To understand whether particular classes of metabolites are more likely to increase or decrease in correlation as a function of health status, we used annotations for each metabolite measurement^19^ (Supplementary Table 1) coupled with hypergeometric tests. Specifically, we asked whether pairs of metabolites significantly affected by health status were more likely to come from each annotation category, relative to the background set of all tested metabolite pairs. We performed these enrichment tests separately for two groups of metabolite pairs, namely (i) those that exhibited stronger correlations in healthy people relative to individuals with metabolic syndrome and (ii) those that showed the opposite pattern. We corrected multiple hypothesis testing using a Bonferroni approach, and report the results in Figure 3 and Supplementary Table 2.

#### Predictive power of metabolite pairs that lose correlation following the onset of disease

Because our dataset included metabolite data collected from the same individuals across multiple time points (sample sizes: n=1564 in 2001, 1498 in 2007, and 1501 in 2011), we asked whether metabolite pairs that became dysregulated (i.e., lost correlation) following the onset of disease could be used to predict which individuals would develop metabolic syndrome at a later time point. To do so, we first identified metabolites with the strongest evidence for dysregulation in our dataset (strongly correlated in healthy people but uncorrelated in those with metabolic syndrome). Specifically, we identified 34 metabolites for which >0.25 of all tested pairs (involving the focal metabolite and any other metabolite) were significantly different between the healthy and metabolic syndrome classes in the expected direction. We performed PCA on these 34 metabolites (using data from first time point), and used the first principal component to predict an individual’s health status at the last time point. To do so, we used a linear model controlling for age and sex at the first time point. For comparison, we performed parallel analyses using the 34 metabolites with the strongest mean differences between the healthy and metabolic syndrome classes.

### The Netherlands study of depression and anxiety (NESDA)

#### Study subjects and sample collection

Our correlation QTL analyses focused on phenotype, genotype, and gene expression data derived from the Netherlands Study of Depression and Anxiety (NESDA; n=5339 participants). NESDA is a previously described cohort study designed to investigate the long-term consequences of depressive and anxiety disorders^15^. Briefly, whole blood was collected in PAXgene Blood RNA tubes from each NESDA participant, and the Affymetrix Genome-Wide Human SNP Array 6.0 and Human Genome U219 Array were used for genotyping and mRNA quantification, respectively. A number of additional health, demographic, and biochemical traits were also recorded for each participant as described in^44^. Complete white blood counts, consisting of lymphocytes, neutrophils, basophils, monocytes, and eosinophils counts were measured for a subset of blood samples.

NESDA contains twin pairs and families discordant for depressive and anxiety disorders, as well as unrelated individuals. To generate a discovery dataset of unrelated individuals for correlation QTL analysis, we removed all members of a given family except for one randomly selected member (leaving us with n=2477 unrelated individuals). For each family, we then randomly assigned one of the remaining individuals to our replication cohort (leaving us with n=1337 individuals).

#### Genotype and gene expression data

Information on genotype and gene expression microarray data generation and quality control are described in detail elsewhere^44^. We filtered the genotype data from our discovery sample set using the following criteria: >5% minor allele frequency (MAF), >5% of individuals exhibit the homozygous minor genotype, and the focal SNP is not in linkage disequilibrium (LD; r^2^<0.5) with other genotyped SNPs within 50kb. LD filtering was performed using the ‘snpgdsLDpruning’ function in the R package *SNPRelate*^45^. This filtering left us with 93197 SNPs for analysis.

Gene expression data were first log2 transformed and filtered to remove probe sets measuring lowly expressed transcripts. Specifically, we calculated the mean expression level in our discovery cohort for every probe set, and excluded probe sets if this value exceeded the maximum value obtained for any control probe set (which should all theoretically have an expression level of 0). Probe sets were also removed if they overlapped a genotyped, polymorphic SNP in the NESDA cohort or a SNP at >5% MAF in whole genome-sequencing data from 769 Dutch individuals^46^. To identify the genomic location of each probe set for this purpose, we downloaded all 25bp probe sequences for the Human Genome U219 array, mapped these sequences with *STAR*^47^ to the human reference genome (hg19), and extracted a set of genomic location coordinates for each probe set from the alignment file. We performed intersections of these genomic location coordinates with known SNP locations (downloaded from the Affymetrix website) using *bedtools* ^48^.

### Testing for genetic effects on variation in correlation

#### Mapping correlation QTL in NESDA

We used the filtered set of SNPs and expression probes measured in NESDA to identify associations between individual SNPs and variation in correlation between the expression levels of two transcripts (referred to here as ‘correlation QTL’). Because our dataset is not well-powered to test all possible pairwise combinations of gene expression probes against all genotyped SNPs, we explored several ways to reduce the search space (i.e., to reduce the number of SNPs and/or genes to test for correlation QTL). Specifically, we attempted analyses focusing on: (i) pairs of known transcription factors and their target genes^21^, tested against SNPs within 1MB of either gene; (ii) pairs of genes known to be involved in the same biological pathway^49^; and (iii) pairs of genes associated with a trait of interest (specifically, BMI, age, or smoking behavior (smoker/non smoker)). In the main text, we report results for analyses that tested for correlation QTL at BMI-associated genes, as this is the approach that produced the strongest signal.

To identify BMI-associated genes, we constructed a linear model for each filtered probe set measured in the NESDA discovery set. Specifically, we tested for an association between log2 transformed expression levels and BMI, controlling for sex, age, smoking behavior (smoker/non smoker), diagnosis with major depressive disorder (yes/no), red blood cell counts, year of sample collection, study phase, and the first 5 principal components from a PCA on the filtered genotype call set (obtained from the ‘prcomp’ function in R). We extracted the p-value associated with the BMI effect from each model, and rank ordered probe sets according to this statistic. We retained the top 300 unique genes for downstream analyses, which corresponded to 475 unique probe sets (because multiple probe sets, in most cases tagging different exons, exist on the array for a given gene).

To calculate correlation among the set of 475 gene expression measurements associated with BMI, we used a slightly modified version of the approach described for our metabolite analyses (see *Comparison of approaches for quantifying correlation*). Specifically, we first removed any mean effects of the following major covariates by running a linear model on the log2 transformed expression values and extracting the residuals: BMI, sex, age, smoking behavior, diagnosis with major depressive disorder, red blood cell counts, year of sample collection, and study phase. We further normalized the residuals for each probe set to mean 0 and unit variance using the ‘scale’ function in R, and computed the product of all possible pairwise probe set combinations (n=112575 combinations, equivalent to 475 chose 2). We used this matrix of products as the input for our QTL analysis. To map genetic effects on variation in correlation, we used *matrixEQTL*^50^ to test for an association between each SNP passing filters (n=93197) and each vector of products, controlling for BMI, sex, age, smoking behavior, diagnosis with major depressive disorder, and the first 5 principal components from a PCA on the filtered genotype call set. We considered a given SNP to be associated with variation in correlation if it passed a 10% FDR threshold^51^.

#### Annotation and analyses of correlation QTL

We identified 484 associations between a given SNP and variation in correlation between two probe sets in our NESDA discovery dataset. For this list of 484 associations, we performed several follow-up analyses to gain biological insight. First, we asked whether the set of genes involved in significant correlation QTL were enriched for (i) known transcription factors or (ii) known targets of transcription factors compared to the background set of all genes tested for correlation QTL. To do so, we used a database of transcription factor-gene target associations and hypergeometric tests^21^.

Second, we asked whether the set of genes involved in significant correlation QTL were enriched for specific gene ontology categories^22^ compared to the background set of all 300 genes tested (Supplementary Table 4). To do so, we used the R package ‘mygene’ and hypergeometric tests.

Third, many of the correlation QTL involved one gene, *RAP1GAP*, whose primary function is to switch the transcription factor *RAP1* from its active GTP-bound form to its inactive GDP-bound form. Therefore, we asked whether genes involved in correlation QTL with RAP1GAP were more likely to be bound by *RAP1*, using ChIP-seq data from mouse embryonic fibroblasts^26^. To do so, we lifted over coordinates for 30398 *RAP1* binding sites from the mouse genome (mm9) to the human genome (hg19) using the UCSC Genome Browser *liftOver* tool^52^. We then tested (using Fisher’s Exact tests) whether genes involved in correlation QTL with *RAP1GAP* were more likely to be bound by *RAP1,* compared to genes not involved in any significant correlation QTL (n=139 genes). We also assigned each gene to its nearest *RAP1* ChIP-seq peak, and tested whether the absolute distance to the nearest peak (from the gene’s annotated transcription start or end site^52^) significantly differed for either of the two comparisons sets described above. If the nearest *RAP1* ChIP-seq peak was within the gene, the distance was coded as 0. We used Wilcoxon signed-rank tests to perform these comparisons.

#### Replication of correlation QTL in NESDA and YFS

For the list of 484 correlation QTL identified in our NESDA discovery set, we performed association tests in the NESDA replication group (n=1337) following the same procedures described in *Mapping correlation QTL in NESDA*.

For each of the 484 correlation QTL, we also implemented parallel association testing in the YFS (n=1414) for probe-SNP combinations that met the following criteria: (i) probes that measured both of the focal genes passed all quality control and expression level filters in YFS and (ii) the focal correlation SNP was also typed in YFS. This filtering criteria left us with 74 associations to potentially replicate in YFS. To do so, we first removed mean effects of BMI, sex, age, smoking behavior, and sampling location using linear models applied to log2 transformed expression values. We normalized these residuals using the ‘scale’ function in R, and computed the product for each of the 74 focal probe pairs. We used these products to test for an effect of each focal SNP, controlling for BMI, sex, age, smoking behavior, and the first 5 principal components from a PCA on the filtered genotype call set.

### Comparison of approaches for quantifying correlation

Our analyses of metabolic syndrome effects on variation in correlation focused on metabolite data that were z-score normalized within each health status group (metabolic syndrome versus healthy). The parallel of this approach for mapping correlation QTL would be to z-score normalize each transcript within each genotypic class before computing the product between two focal transcripts. Such an approach has the advantage of removing any mean or variance effects of the predictor variable (e.g., health status or genotype) on the outcome variables (e.g., metabolite or gene expression levels). However, this specific pipeline is infeasible for genome-wide QTL mapping, as it would require us to recalculate the outcome variable for every association test we performed (i.e., we would need to re-normalize each transcript pair and re-calculate the product every time we tested a new SNP for an association with correlation). This would be extremely computationally costly, and would preclude us from using the efficient tools for QTL mapping that made our genome-wide genetic screen feasible^50^.

As an alternative, we therefore regressed out mean effects of many major covariates from our gene expression data, and normalized the resulting residuals, before computing the product between two transcripts. This approach is different than normalizing each transcript within each genotypic class, but attempts to circumvent the same set of potential issues and has the advantage of only needing to be performed once. For comparison, we present the effects size estimates and p-values from the two approaches for the set of 484 significant correlation QTL we identified (Supplementary Figure 6).

### Cell type heterogeneity-related confounds

Our correlation QTL analyses focused on gene expression levels measured in whole blood, which is composed of several cell types with distinct transcription profiles. All of our analyses conducted with NESDA data controlled for one measure of cell type heterogeneity – the proportion of red blood cells in a given sample. However, this crude measure may not capture the full spectrum of potential confounding effects of cell type heterogeneity. In particular, we consider two possible scenarios: (i) BMI is associated with finer-grained measures of cell type heterogeneity, such that the genes we tested for correlation QTL are actually only associated with cell type composition (and not BMI) and (ii) genotypes at the correlation QTL we identified are associated with finer-grained measures of cell type heterogeneity and thus produce a false positive signal. Under this scenario, individuals of a given genotype would have high expression levels of two focal genes (and thus high values of their product) because of variation in cell composition, rather than genotypic effects on correlation. We note that scenario (i) would complicate the biological interpretation of our results, but would not produce the strong type I error expected under scenario (ii).

To investigate the evidence for scenario (i), we used data on the number of eosinophils, basophils, neutrophils, lymphocytes, and monocytes in each whole blood sample, which were available for a subset of our NESDA discovery set (n=594). We used these data to test (using linear models) for an association between each measure of cell type composition and BMI, controlling for the covariates we included in all analyses (namely, sex, age, smoking behavior, diagnosis with major depressive disorder, red blood cell counts, year of sample collection, and study phase). We found weak or no associations between most cell measures and BMI (p<0.01, percent variance explained by BMI<1%) except neutrophil composition. Here, BMI explained 4.7% of the variance (p=1.88×10^-8^). We note that these modest effects may indicate that our set of BMI-associated genes are somewhat ‘noisy’, but given the magnitude of the BMI-neutrophil association we expect the impact of these effects on our results to be minimal.

Under scenario (ii), we would expect genotype to be significantly associated with the mean expression levels of the two focal genes involved in the correlation QTL. We note that we have already explored the contribution of this scenario to the signal we observe (see *Simulations*; Table 1 and Supplementary Figure 1).

## Acknowledgments

We thank all volunteers who participated in the studies described here. We also thank the Ayroles lab for helpful comments on previous manuscript versions. This study was funded by National Institutes of Health (NIH) grants GM124881 to JFA, HL-095056 and HL-28481. AJL is supported by a postdoctoral fellowship from the Helen Hay Whitney Foundation, and AK is supported by NIH grant F31HL127921. The Young Finns Study has been financially supported by the Academy of Finland: grants 286284, 134309 (Eye), 126925, 121584, 124282, 129378 (Salve), 117787 (Gendi), and 41071 (Skidi); the Social Insurance Institution of Finland; Competitive State Research Financing of the Expert Responsibility area of Kuopio, Tampere and Turku University Hospitals (grant X51001); Juho Vainio Foundation; Paavo Nurmi Foundation; Finnish Foundation for Cardiovascular Research; Finnish Cultural Foundation; Tampere Tuberculosis Foundation; Emil Aaltonen Foundation; Yrjö Jahnsson Foundation; Signe and Ane Gyllenberg Foundation; Diabetes Research Foundation of Finnish Diabetes Association; and EU Horizon 2020 (grant 755320 for TAXINOMISIS).

## Data and code availability

The NESDA dataset was accessed through dbGaP (study accession: phs000486.v1.p1). The YFS dataset is available to the scientific community based on a written application, and further information can be found at www.utu.fi/med/cardio/youngfinnsstudy. Code for implementing the test for differential correlation we describe here can be found at https://github.com/AmandaJLea/differential_correlation.

